# How far can microbial monocultures predict growth in multi-strain communities?

**DOI:** 10.64898/2026.09.02.748814

**Authors:** Florian Borse, Silvana Lord Smits, T. Anthony Sun, Johannes Cairns, Anne Farewell, Ville Mustonen

**Affiliations:** Department of Organismal and Evolutionary Biology, Department of Computer Science, University of Helsinki, 00014 Helsinki, Finland; Department of Chemistry and Molecular Biology, University of Gothenburg, 41390 Göteborg, Sweden; Center for Antibiotic Resistance Research in Gothenburg (CARe), 40530 Göteborg, Sweden; Department of Experimental Medical Science, Lund University, 221 84 Lund, Sweden; Science for Life Laboratory (SciLifeLab), Lund University, 221 84 Lund, Sweden; Turku Collegium for Science, Medicine and Technology, and Department of Biology, University of Turku, 20014 Turku, Finland

## Abstract

Previous research in microbes successfully predicted biculture growth based on monoculture growth curves. Still, the usual model-based approach does not seem to extend to communities involving more than two bacterial strains. Here, we use a model-blind machine-learning approach to predict community-wide yield, area under the growth curve, maximum relative growth rate and its timing in communities involving up to five strains of *Escherichia coli*. First, we identify the highest per-capita growth rate and its timing in monocultures as major predictors of multi-strain community growth. Next, we show that a random forest trained on communities involving a low number of strains is able to predict the outcomes of communities involving a higher number of strains. Finally, we reveal diminishing returns in using more and more complex communities in order to predict the behaviour of a higher-level community, because monoculture- and biculture-based growth predictions are often already accurate. This finding relativises the need for experiments involving high numbers of strains when studying the growth dynamics of multi-strain communities.

**Author summary:** It is possible to predict how two bacterial species grow when we culture them together, based on how each species grows in isolation. However, common approaches do not work when more species are cultured together. In this study, we use machine learning methods to study how the predictability of collective growth behaves, according to patterns and aspects learned from the growth of subcommunities in isolation. We show that community-wide growth is significantly impacted by which species reaches the earliest a critical per-capita growth rate when introduced in a fresh culture medium. We also find that most of the predictability stems from knowing single-species behaviour as well as species pairs. This result suggests that it is not necessary to investigate all intermediate subcommunity levels to predict how the full community grows.

## Introduction

Recent microbiological research has identified the key role played by the multispecies context in shaping a bacterial community’s diversity [1], metabolic niche [2] and resistance to antibiotics [3], with paramount implications for human health [4]. In that context, it has become an essential problem in current microbiology to quantify higher-order interactions (HOIs), that is emergent properties [5, 6] by which community-level behaviours do not simply result from phenomena at the level of single species or of species pairs [7, 8]. Indeed, whether pairwise interactions completely capture interspecific interactions or whether, on the contrary, involving more species generally gives rise to HOIs [9, 10] is often unclear, system- and context-dependent [11, 12], but has an enormous impact on our ability to predict the growth of a multi-strain community [13].

Usually, assessing HOIs relies on a logistic-based model, such as the generalised Lotka-Volterra model [12, 14, 15]. This mathematically simple approach yields good fits in bicultures starting from two species introduced in equal proportions into fresh culture medium, on the basis of monoculture-inferred parameters [16], and readily extends to multiple interacting species in theory [17]. However, predictions seem incompatible with observations in multispecies communities [18–22] and to generally depend on empirical conditions [23]. Recent modelling efforts have thus attempted to account for key biological mechanisms such as resource competition [20, 24–27].

Predicting the behaviour of complex communities has relied on formulating an explicit mathematical model for microbial growth. But a model-independent method has been missing, to assess how monocultures inform us about the phenotypic growth properties of multispecies and multi-strain communities. However, model-blind approaches relying on machine learning algorithms are now able to accurately account for microbial growth in experiments, when boosted with biological insights [28, 29]. Although such algorithms are usually black boxes, their inherent properties can be computed, such as the respective importances of input variables, which can in turn inform both experiments and our mathematical modelling arsenal [30].

Here, we assess the extent to which single-strain cultures inform the growth of multi-strain communities. Namely, we obtain growth curves from five strains of *E. coli* cultured in all possible combinations, *i*.*e*., as monocultures, bicultures, tricultures, up to the 5-strain community, across 7 environments (Figure 1). Such combinatorially complete experiments are increasingly feasible and open up new directions for theory development for microbial ecology ranging, for example, from leveraging concepts of cooperative game theory to quantitative genetics [31, 32]. We evaluate the information provided by simpler communities about more complex communities through random forest models’ predictive power, and interpret it biologically using the importances of input variables.

**Fig 1.**
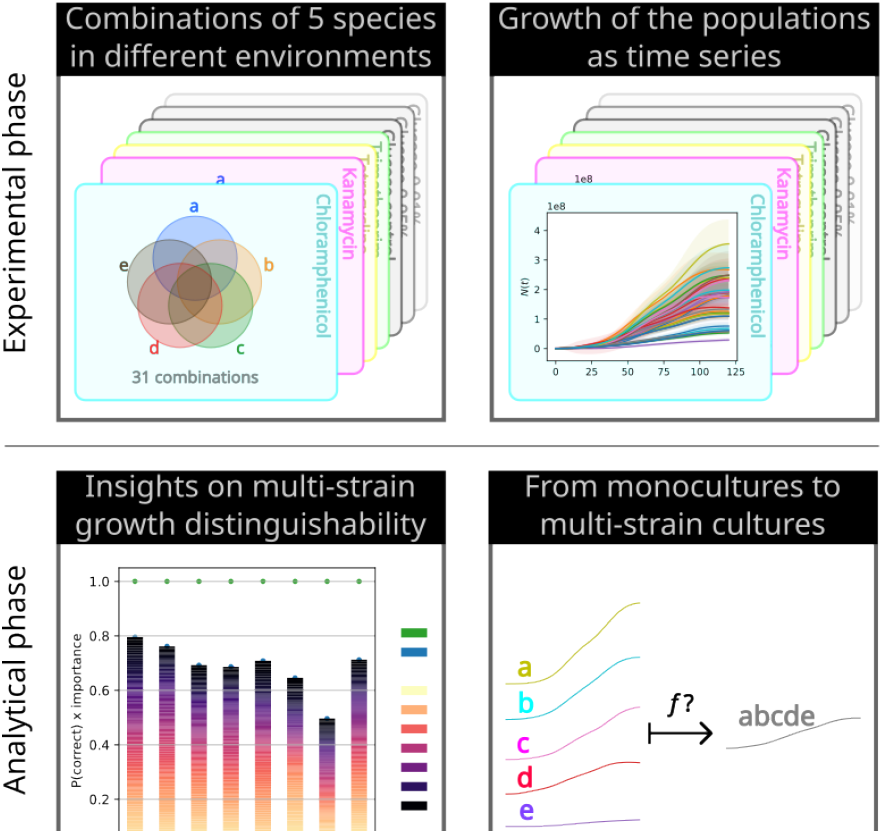
Schematic overview of the experiments and workflow. We first obtain an original dataset where five strains of *E. coli*, labelled as *a, b, c, d*, and *e*, are combined as multi-strain communities for all 31 combinations possible. These communities are grown in seven different environments, and we obtain time series of their growth. We can thus run machine learning algorithms, from which we can gain insight on multi-strain community growth. We start by studying which growth stage random forests would deem as most relevant for distinguishing growth curves of communities of different composition. We then perform regressions from characteristics of monoculture growth to characteristics of growth of multi-strain communities.

First, we identify which stage of growth displays the most characteristic traits distinguishing between combinations of strains and find that early growth bears the most importance for predicting the outcomes of multi-strain communities. Then, we focus on the yield, area under the curve (AUC), maximum growth rate and its timing, which summarise the shape of the growth curves, and show that the summary statistics obtained from monocultures can predict the summary statistics in multi-strain communities reasonably well. Finally, we use partial learning in order to include interactions between strains beyond monoculture information. By iteratively increasing the number of strains in the communities on which the random forests are trained, we successively learn interactions of increasing order and assess their contribution to the models’ performance in predicting the outcomes of arbitrary multi-strain communities. This reveals diminishing returns in predictive power as HOIs are added to the regression models and suggests weak evidence for HOIs when studying growth characteristics.

## Results

### Strain combinations are distinguishable

Growth curves obtained for the same strain combination were considered biological replicates (Figure 2). To assess how much information a growth curve contains, that is how distinguishable different strain compositions’ growth patterns are, we defined two classification tasks.

**Fig 2.**
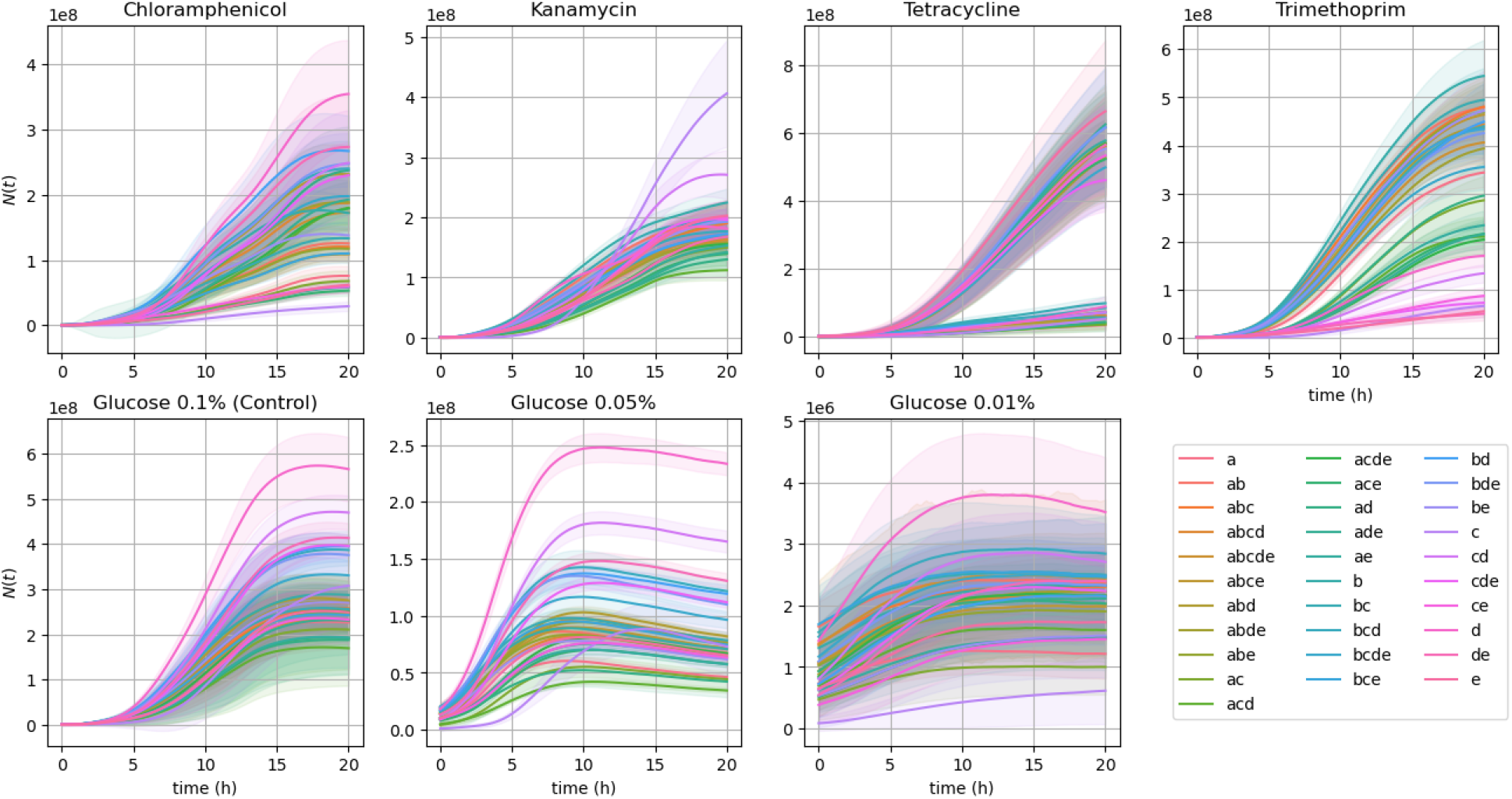
Growth curves (mean ± sd) by environment. Each panel contains population growth curves for a specific environment. Each curve represents the population size *N* (*t*) at a given time *t*, averaged by strain combination (102 curves each, for 120 time points). The curves are thus coloured by strain combination. Additionally, every curve is surrounded by the standard deviation of their strain combination.

The first task consisted of multi-label classifications where, for each environment, we predicted strain composition, based on a growth curve *N* (*t*) given as input. This evaluated whether populations growth curves are distinguishable from each other, and at which growth stage.

The second task consisted of a series of single-label classifications. Each classification predicted, for a given environment, whether a population corresponded to a given strain combination, based on its growth curve *N* (*t*). This assessed the role of combination-specific properties in temporal behaviour and distinguishability.

On average, random forests classified population growth curves by strain combination with 71 % accuracy (Figures 3 and S2). This demonstrated that growth curves contained a substantial amount of information on their strain compositions. Additionally, feature importances showed that earlier time points were generally more informative than later time points, which suggested that the effect of strain composition was stronger during early growth phases.

**Fig 3.**
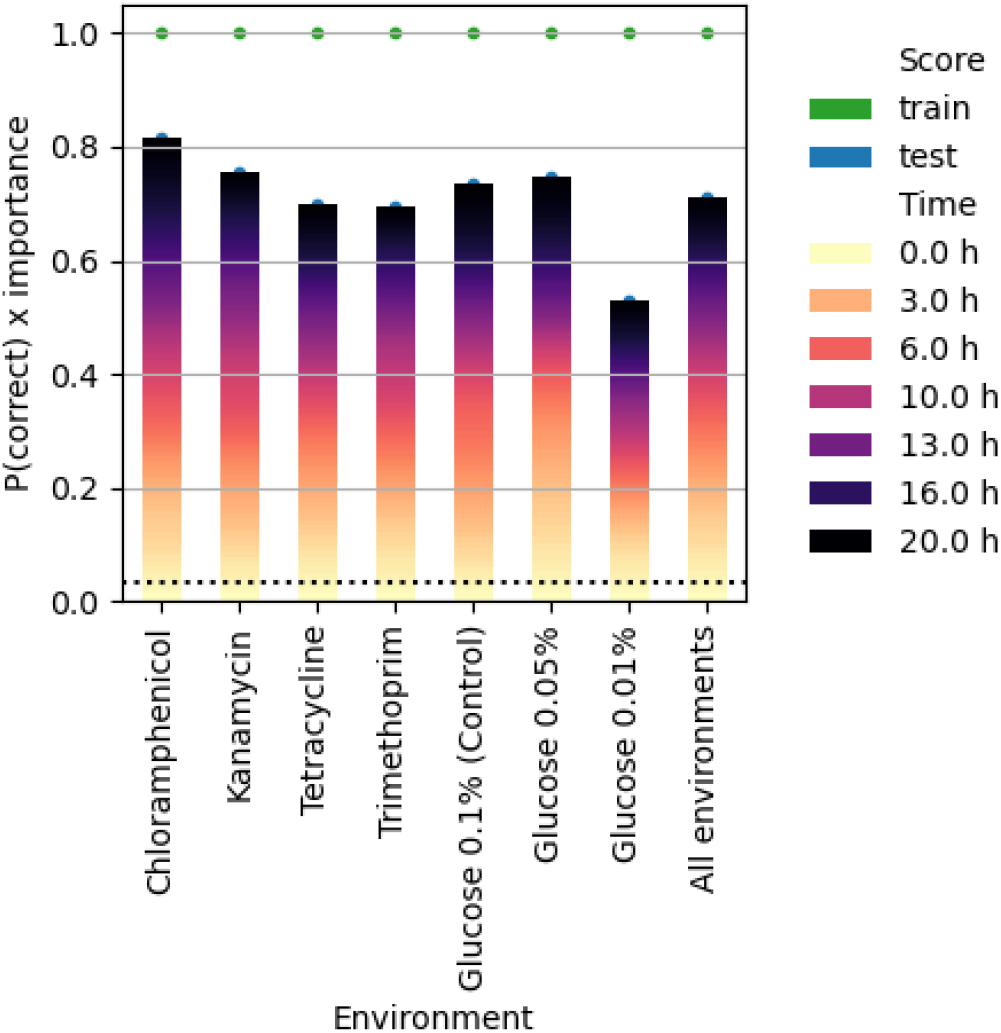
Multi-label classification tasks. Each population growth curve is assessed for whether it can be classified by its community composition. Each bar represents an environment, for which the classifications take place. The bars are coloured according to the importance of a given time for predicting the correct label, i.e., the shorter the dark part of a bar is, the more growth information is carried by the initial stages of growth. The dashed line represents the probability to guess correctly (1/31).

### Multi-strain community growth properties are predictable

Next, we asked whether the collective growth properties of a community are predictable when knowing the growth properties of each member strain in a monoculture. We formulated this problem in terms of regression tasks based on summary values characterising a growth curve. For example, we started by limiting the problem to predicting the growth properties of a biculture *ij* from the properties of monocultures *i* and *j*.

We could then train a random forest on that regression, and assess its performances by using the growth properties calculated for the combination.

To that aim, we used properties commonly defined in population growth models (Figure S1). Namely, assuming a population growth curve *N* (*t*), we calculated its relative growth rates as 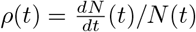, then focused on the following shape parameters: their yield *Y* = *N* (*t*_final_), their maximum relative growth rate *ρ*_max_ = max_*t*_ *ρ*(*t*) and the time until this maximum is reached *τ*_max_ = argmax_*t*_ *ρ*(*t*). We also defined a shape-independent parameter 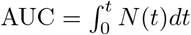.

Writing the growth properties of a monoculture *i* as *S*_*i*_ = (*Y*_*i*_, AUC_*i*_, *ρ*_*i*_, *τ*_*i*_), we could thus write the above regression task in shorter notation as: *S*_*i*_ *S*_*j*_ *↦ S*_*ij*_. This regression problem could be extended to tricultures and above: are the properties of three (or more) monocultures sufficient to predict the growth properties of their combination? *I*.*e*., can a random forest learn the following regression: *S*_*i*_ *S*_*j*_ *S*_*k*_ *↦ S*_*ijk*_?

A variation of this last regression consists in the following multi-strain prediction problem: to predict the behaviour of any multi-strain community from the parameters of its monocultures. Given arbitrary numbers of *S*_*i*_ as input, we choose to include the parameters of all monocultures, but then for each strain *a, b, · · ·, e* replace the parameters with zero if that strain is not part of the target multi-strain community. Denoting the presence or absence of a strain*i* in a multi-strain community *I* as *w*_*i*_ = 1 if *i ∈ I* or *w*_*i*_ = 0 otherwise, we can thus write this regression as: ⋃_*i*_(*w*_*i*_*S*_*i*_) = *w*_*a*_*S*_*a*_ *w*_*b*_*S*_*b*_ … *w*_*e*_*S*_*e*_ *↦ S*_*I*_.

In most cases, a random forest was able to capture the interactions between two strains (Figure 4, top), although the scores of certain random forests were severely affected by experimental outliers, as the case of *ρ*_max_ in Chloramphenicol (Figure S4). Overall however, knowing how monocultures grew allowed to predict the growth characteristics with 87 % accuracy in bicultures (Figure 4a), 88 % in tricultures (Figure 4b), and 84 % for all multi-strain communities together (Figure 5).

**Fig 4.**
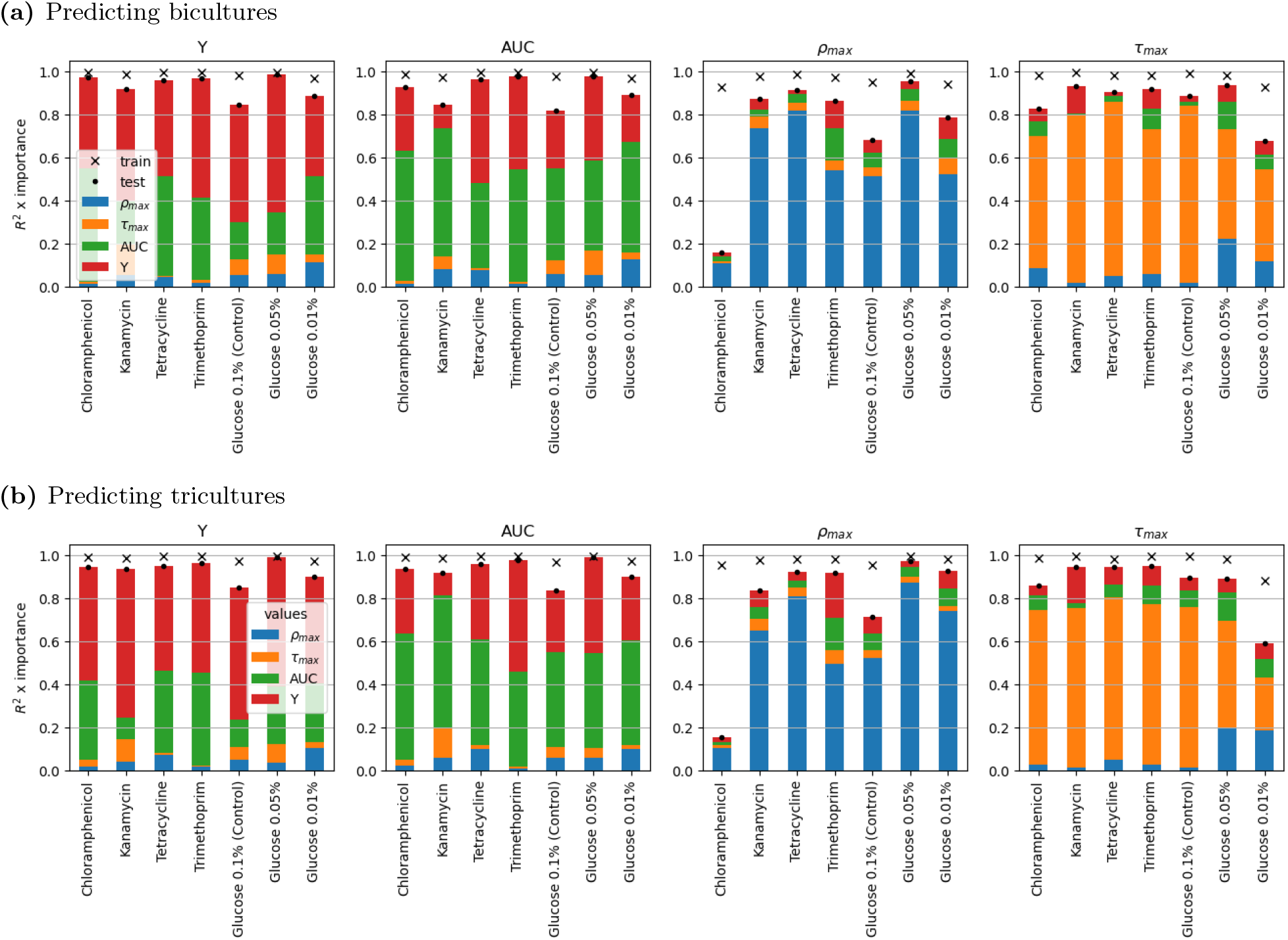
Predicting bicultures and tricultures from monocultures. A random forest is trained to predict summary values for the shape of the growth curves of bicultures and tricultures, using the summary values of the involved monocultures as input. Each panel represents the prediction scores and importance of input features for a specific growth summary variable. Each model is trained on train data, which is three quarters of the data of a specific environment, and then scored (*R*^2^) on the train data and test data, which are displayed respectively as a cross and a dot. Each bar then represents the importances of the input features of a random forest, grouped by growth summary variables. These importances are multiplied by the test score.

**Fig 5.**
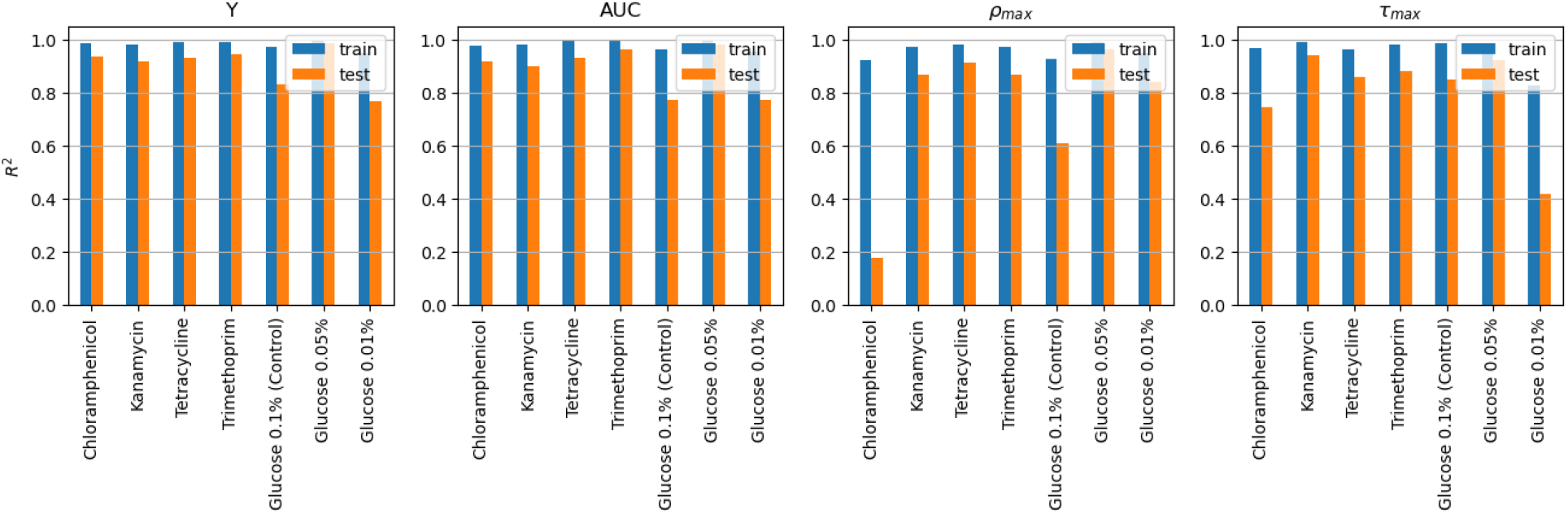
Predicting multi-strain community growth from monocultures. A random forest is trained for each environment and summary value for a growth curve shape, to predict the summary value of any community by using the summary values of all monocultures as input, each set to zero if a strain is not part of the community. The data is split in train and test set the same way as for figure 4.

Feature importances revealed that the final yield and the area under the curve seemed to compete in a linear race to explain the same outcomes. Additionally, the impact of early growth stages uncovered through Figure 3 – here through maximum growth *ρ*_max_ and its timing *τ*_max_ – was not observed for the timing-independent variables *Y* and AUC. A slight variant of these regressions, however, consisting of predictions using only information on *ρ*_max_ and *τ*_max_ to predict *Y* and AUC, allowed to explain this inconsistency. Indeed, as the results of these regressions (Figure S5 showed, the former values allow toed predict relatively well the latter values. This suggests that the importance of *Y* and AUC found in Figure 4 occurred mostly by the regression model favouring directly related input parameters (Y and AUC), as opposed to indirectly related parameters (*ρ*_max_ and *τ*_max_). In practice, we recovered in Figure S5 the impact of early growth stages observed in 3. Extending these regressions successively to four-strain then five-strain communities resulted in similar considerations (Figure S3).

### Information contained in higher-order interactions

The previous regression tasks showed that, by themselves, monocultures contained considerable information to predict the growth of multi-strain communities. However, our focus on monocultures missed an essential component of multi-strain communities: the potential interactions between strains, including delineating possible HOIs undetectable in pairwise experimental contexts.

To find about these interactions, we now assessed the predictability of the same summary values of growth when training models on communities, using member strain presence/absence vectors as input features, with an iteratively increasing number of strains. If one random forest model is trained on communities involving up to *M* strains, and another model is trained on communities involving up to *M −* 1 strains, how much better can the first model predict community-level properties of an arbitrary number *N* of strains? In other words, how much better can a model trained to include interactions up to a certain order perform, compared to a model trained to include interactions up to a lower order?

We devised the following regressions: for each number of strains *M*, a random forest was trained to predict community summary values, given the environment and the strain combinations as input feautures, then tested on predicting summary values for communities combining any higher number *N > M* of strains. Monocultures alone showed relevant predictive power for the maximum growth rate only (Figure 6c). For most growth summary statistics, the highest increase in predictive power is achieved by adding bicultures to the training set. Comparatively, subsequent additions did not allow for substantial improvements, except in the case of the timing of maximum growth (Figure 6d).

**Fig 6.**
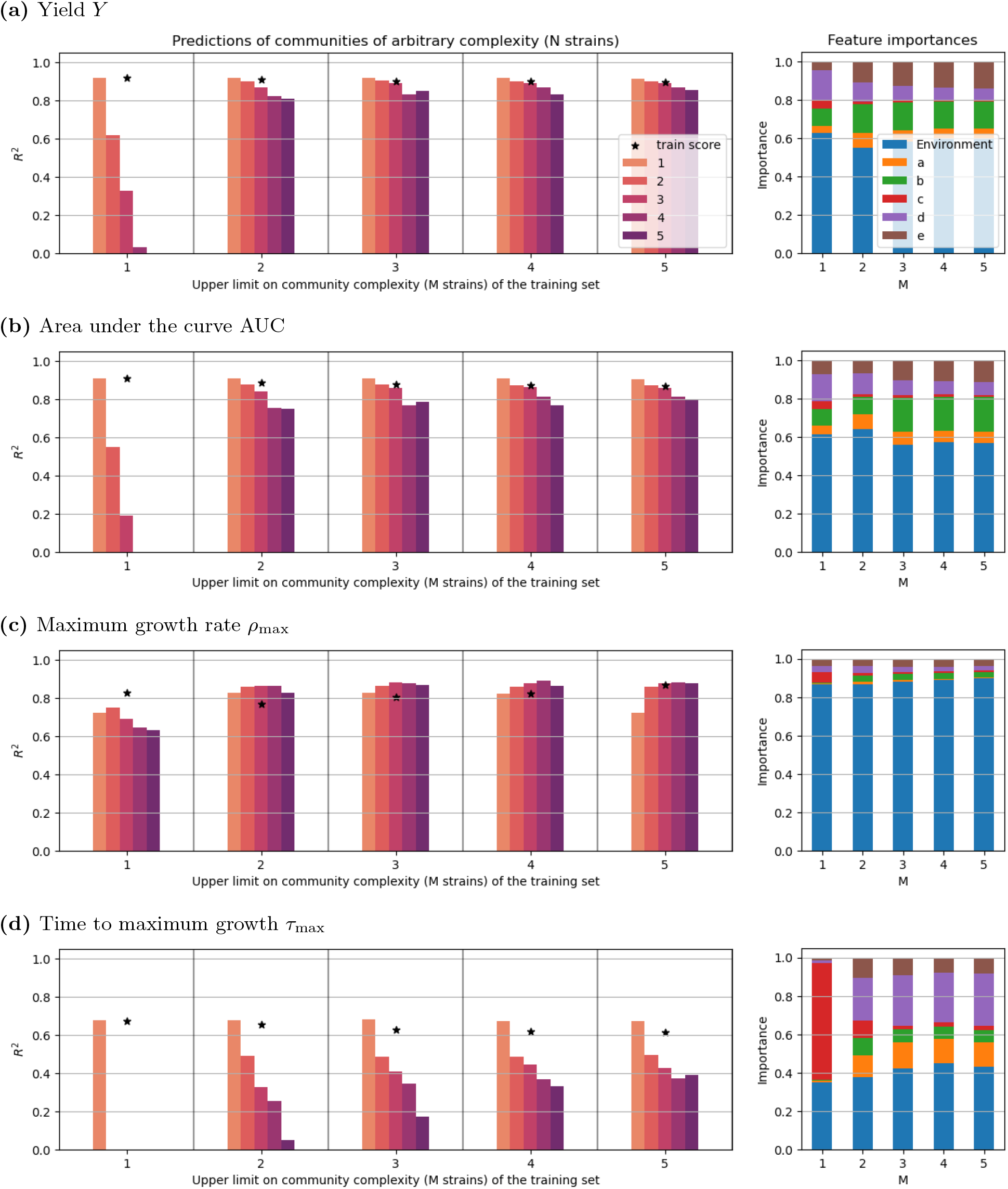
Partial training performances. Random forests have been trained on communities of an iteratively increasing number of strains (*M*), to predict the growth summary values of a multi-strain community given strain presence and environment. The models have then been tested on communities of arbitrary complexity *N*. We also display the feature importances of the random forests for each number *M* of strains considered for training.

This suggests that while first-order interactions between strains contain valuable information, higher order interactions may only bring marginally significant information, at least in our dataset. In other words, monoculture behaviours and pairwise interactions together were sufficient to predict the growth behaviour of more complex strain combinations.

## Discussion

We studied the growth phenotype of five strains of *E. coli* in monocultures and in all of their combinations up to the whole 5-community, under different environmental conditions.

Random forests were able to classify population growth curves by strain combination with 71 % accuracy. We found that earlier stages of growth played a greater role in classifying multi-strain community compositions, compared to later growth (Figures 3 and S2). On the one hand, the comparative importance of the exponential growth phase is expected, as opposed to the stationary phase of microbial growth, in determining the behaviour of the whole community. On the other hand, this result suggests that interactions between strains happen primarily during early community growth. This is in line with the idea that they are mainly mediated by resource competition, as a growing body of studies shows [20, 24–27, 29].

Next, we focused on the yield, area under the curve, maximum growth rate and its timing, in order to summarise growth curves and perform regression tasks. Monoculture information yielded fairly good predictions overall of the growth characteristics for multi-strain communities. We identified the maximum per-capita growth rate *ρ*_max_ and its timing *τ*_max_ as crucial monoculture growth features to predict multi-strain cultures (Figure S5). This pattern explains how communities may be distinguished by their strain composition, through a mechanism combining the specific values for *ρ*_max_ and *τ*_max_ into community-wide, collective growth rate and timing. Similar to the early-bird effect observed in microbial community assembly [33], we propose the null-hypothesis that these collective *ρ*_max_ and *τ*_max_ mainly result from the strain which reaches a critical growth the earliest, thus consuming the resources necessary for growth. Such priority effects are widespread in community ecology [34, 35] and could result in resource monopolisation, with the potential to crucially impact the community’s eco-evolutionary dynamics [36, 37].

Compared to training our models on monocultures only, we found that training on both monocultures and bicultures substantially improved prediction accuracy for all growth characteristics. However, including information on 3-strain and 4-strain communities did not yield any further substantial improvement. This suggests that strain growth characteristics and pairwise interactions observed in monocultures and bicultures effectively capture most of the information on multi-strain growth [38]. In line with other experimental studies [5, 39, 40], this finding challenges the idea that HOIs play a major role in the macroscopic growth behaviour of microbial communities.

This result concerns the whole community’s macroscopic growth specifically, and remains compatible with HOIs discussed when studying other community-level properties such as stability, assembly and metabolic functions [18, 20, 21, 41–43], community composition and species coexistence [44–46], or the collective capacity to detoxify antibiotics [47]. It is worth noting that HOIs are generally also motivated by theoretical expectations [8, 48] in general community ecology, with similarly unclear evidence for their actual role in explaining empirical observations [9, 13, 49]. Additionally, HOIs could also play a greater role in the case of phylogenetically more distant organisms and/or in the context of a more complex culture medium, as opposed to our simple experimental setting involving five *E. coli* strains in minimal medium. Further research is required to extend or limit our conclusions to quantify how close the behaviour of our multi-strain collective is to a single-species dynamics.

## Materials and Methods

### Experimental protocols

#### Strains and strain mixtures

Five different *E. coli* isolates were used in this experiment. These strains were previously selected by Gamfeldt et al. (2023) [50] from a large-scale screening of 768 isolates, exposed to sub-minimum inhibitory concentration (sub-MIC) levels of various antibiotics. The selection was based on their modestly different growth yields under various conditions (tetracycline, chloramphenicol, trimethoprim, kanamycin). And we used the following shorthand names for the different isolates [50]:

a GU1715

b GU1910

c GU1969

d GU2134

e GU2295.

Throughout the project, *E. coli* ATCC 25922, commonly used as a control when conducting antibiotic susceptibility tests, is used as a spatial control.

Isolates were cultured overnight in LB Miller broth (10 g/L NaCl) at 37 degree Celsius with shaking (180 rpm). All cultures were subsequently adjusted with fresh LB medium so that their optical densities at 600 nm (OD600) reached 0.5. The adjusted cultures were mixed in all possible combinations of up to 5 strains, with equal proportions of each isolate in the final mixtures (i.e., two strains were mixed 1:1 based on OD and 5 strains were mixed in the ratio of 1/5 each). The resulting mixtures were diluted with glycerol to a final concentration of 15 % glycerol to allow for freezing. 200 microlitres of all mixtures were then added to different wells of a 96-well pate in a randomised format, as previously done by Gamfeldt et al. (2023) [50], and as shown below (Table 1). Bordering wells on the plate were filled with ATCC 25922 in 15% glycerol to avoid edge-effects due to greater nutrient availability on edges in downstream experiments on agar plates. A second 96-well plate was prepared with ATCC 25922 and 15% glycerol in every well, later used as a spatial control. Once prepared, the 96-well plates were stored at −80 degrees Celsius until use.

**Table 1.** 96-well plate lay-out of strain mixtures.

|  | 1 | 2 | 3 | 4 | 5 | 6 | 7 | 8 | 9 | 10 | 11 | 12 |
| --- | --- | --- | --- | --- | --- | --- | --- | --- | --- | --- | --- | --- |
| A | ATCC | ATCC | ATCC | ATCC | ATCC | ATCC | ATCC | ATCC | ATCC | ATCC | ATCC | ATCC |
| B | ATCC | de | b | ae | abe | bc | ac | abce | abcde | bce | abc | ATCC |
| C | ATCC | abde | bd | ce | e | ace | acde | cd | d | ad | ab | ATCC |
| D | ATCC | bcde | be | abcd | de | abd | b | ae | abe | bc | c | ATCC |
| E | ATCC | bcd | ac | abce | cde | abcde | acd | bce | bcde | ab | bde | ATCC |
| F | ATCC | ade | c | bde | a | abde | abc | bcd | ade | acd | a | ATCC |
| G | ATCC | bd | ce | be | ace | acde | cd | d | ad | e | abcd | ATCC |
| H | ATCC | ATCC | ATCC | ATCC | ATCC | ATCC | ATCC | ATCC | ATCC | ATCC | ATCC | ATCC |

#### Media

M9 minimal medium [51] (0.01 mM thiamine, 15 g/L agar and 0.1% glucose) was used unless specified. It was used to generate all preculture plates. The test plates we used were M9 plates containing three concentrations of glucose (0.01%, 0.05%, or 0.1%) and M9 plates with 0.1% glucose and sub-MIC concentrations of each of the four antibiotics (1*µg/ml* chloramphenicol, 8*µg/ml* kanamycin, 0.3*µg/ml* tetracycline and 0.25*µg/ml* trimethoprim).

#### Pinning

To generate preculture plates, both the sample plate, containing strain mixtures, and the control plate, containing only ATCC 25922, were used. After thawing, they were jointly pinned onto agar media (0,1% glucose preculture plates) using an HDA RoToR robot (Singer LTD, UK) to a density of 1536 colonies per plate. Every fourth position on these plates originated from the control plate, all other positions originated from the sample plate. Thus, every well on the sample plate was pinned 12 times onto each preculture plate, and each well on the control plate was pinned 4 times per preculture plate. A separate preculture plate was used per test plate. After pinning, plates were incubated at 30 degrees Celsius for 16 hours. The HDA RoToR robot (Singer LTD, UK) was then used to pin all colonies from each preculture plate to a test plate, containing control media or one of the various selective conditions. Two test plates per condition were included. Test plates were subsequently placed in Epson Perfection V800 photo scanners (Epson Corporation, United Kingdom), kept in a humidity- and temperature-controlled cabinet at 30 degrees Celsius, and incubated for 48 hours.

The Scan-o-matic software [52] was used to image the test plates every 10 minutes during the 48 hours. Through use of this software, obtained pixel intensities for each colony were transformed into population size measures (calibrated for *E. coli* [53]).

### Preprocessing of the data

The data was normalized to the fourth position controls, and raw growth curves were obtained for each position. Several scanners were involved for the data acquisition. Distributing each environment across at least two different scanners brings the advantage of guaranteeing that scanner failure does not affect all copies of an experiment. This means also that the first step is to bring all the data under a common uniform pool.

Since all experimental cultures reached stationary phase by time point 120, we retained only the first 120 time points (20 hours). As derivatives – *ρ*_*i*_(*t*) in particular – are considered, the data is also first interpolated and then smoothened using an averaging window of 10 time points.

Not every growth curve can be interpolated, mainly due to missing initial time point if there are any data at all, which means that the curve data needs to be removed. Moreover, some growth curves display severe technical artefact characteristics; to eliminate these, we apply a per plate cut-off based on the initial population size. Finally, due to spatial constraints on a 96-well plate, two patches of replicas for each combination of strains cannot be guaranteed, and two combinations of strains do indeed only have one patch per plate.

This creates an inconsistency in number of replicas per environment per strain combination. As some of our operations require consistency, and as the number of available curve replicas is otherwise high enough, we choose to bootstrap the eventual missing growth curves by simply cycling through the available curves once. This ensures that a certain curve replica is present at maximum two times.

### Random forest models

For random forest implementations, we used the sklearn.ensemble module’s RandomForestClassifier and RandomForestRegressor classes from the scikit-learn library [54], with default hyperparameters.

We split the data into train and test sets by taking for every kind of data each third sample as training sample, and using the others as testing samples. Since each kind of data was distributed as 4 × 4 patches, taking each third sample allowed to avoid spatial biases in the sampling, regardless of the ML task.

Classification tasks were evaluated by comparing the predictions for the test set with the test set data, and calculating the probability as number of correct labels divided by total number of test samples.

Regression tasks were evaluated using the built-in implementation of the *R*^2^ score, for the predictions for the test set compared to the test set data. The random forest implementations from scikit-learn provide a feature importances method [55], which calculates the contribution of each model variable to future predictions based on mean decrease in impurity.

## Code and data availability

All code and data in support of this publication are available at https://doi.org/10.17605/OSF.IO/NSR9H.

## Author contributions

**Conceptualization:** Florian Borse, Anthony Sun, Anne Farewell, Ville Mustonen

**Formal analysis:** Florian Borse, Anthony Sun, Ville Mustonen

**Investigation:** Florian Borse, Silvana Smits, Anthony Sun, Johannes Cairns, Anne Farewell, Ville Mustonen

**Methodology:** Florian Borse, Anthony Sun, Ville Mustonen

**Software:** Florian Borse

**Experiments:** Silvana Smits, Anne Farewell

**Supervision:** Anthony Sun, Anne Farewell, Ville Mustonen

**Writing – original draft:** Florian Borse, Anthony Sun, Silvana Smits

**Writing – review & editing:** Florian Borse, Anthony Sun, Silvana Smits, Johannes Cairns, Anne Farewell, Ville Mustonen

## SI figures Acknowledgments

The authors wish to thank CSC — IT Center for Science, Finland, for computational resources.

---

**Fig S1.**
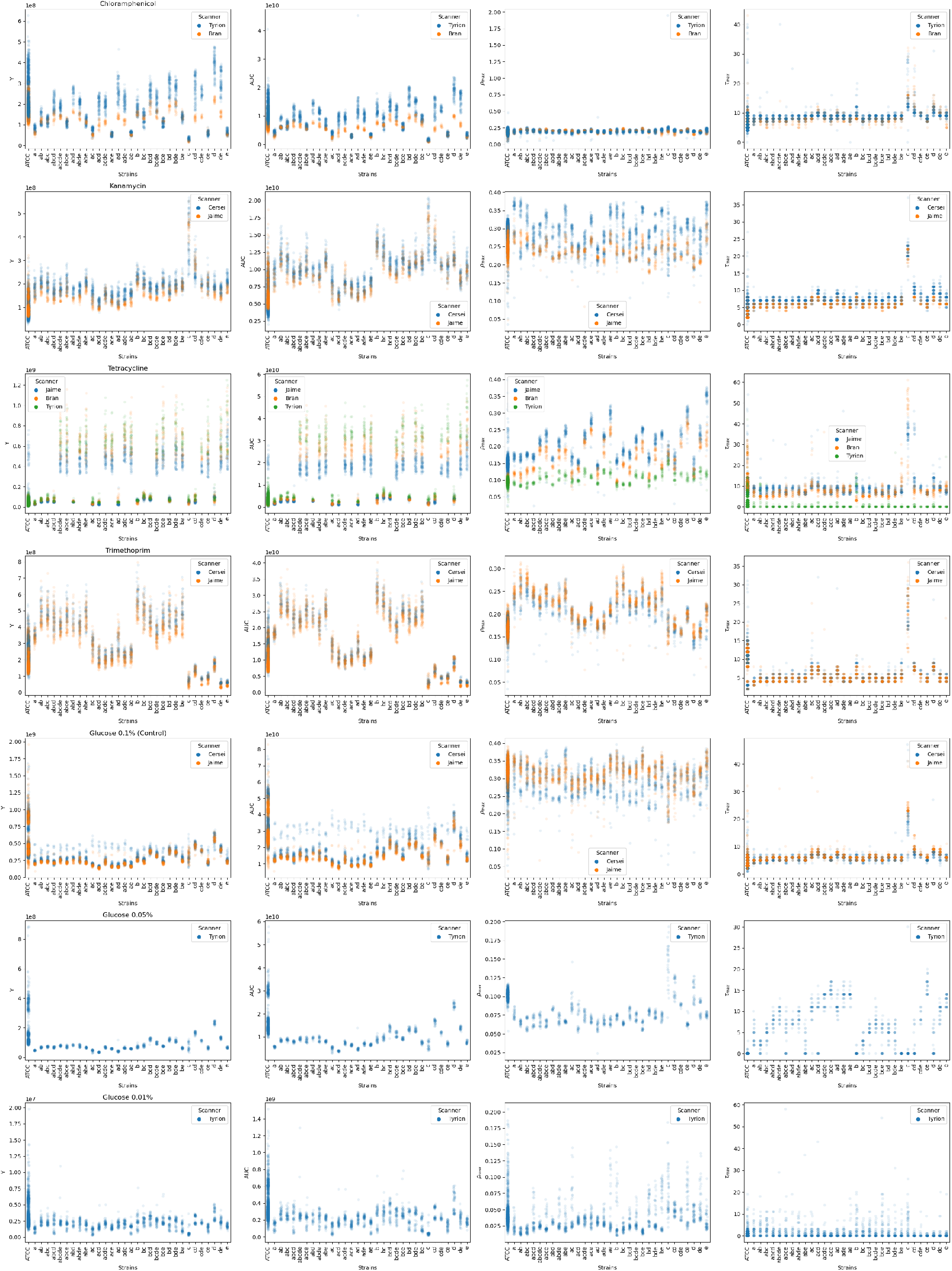
Homogeneity of the summary values. For each population, we compute the following shape values: the yield *Y*, the maximum growth rate *ρ*_max_, its timing *τ*_max_. Additionally, we compute the area under the curve AUC. We then display for each environment and multi-strain composition separately the values.

**Fig S2.**
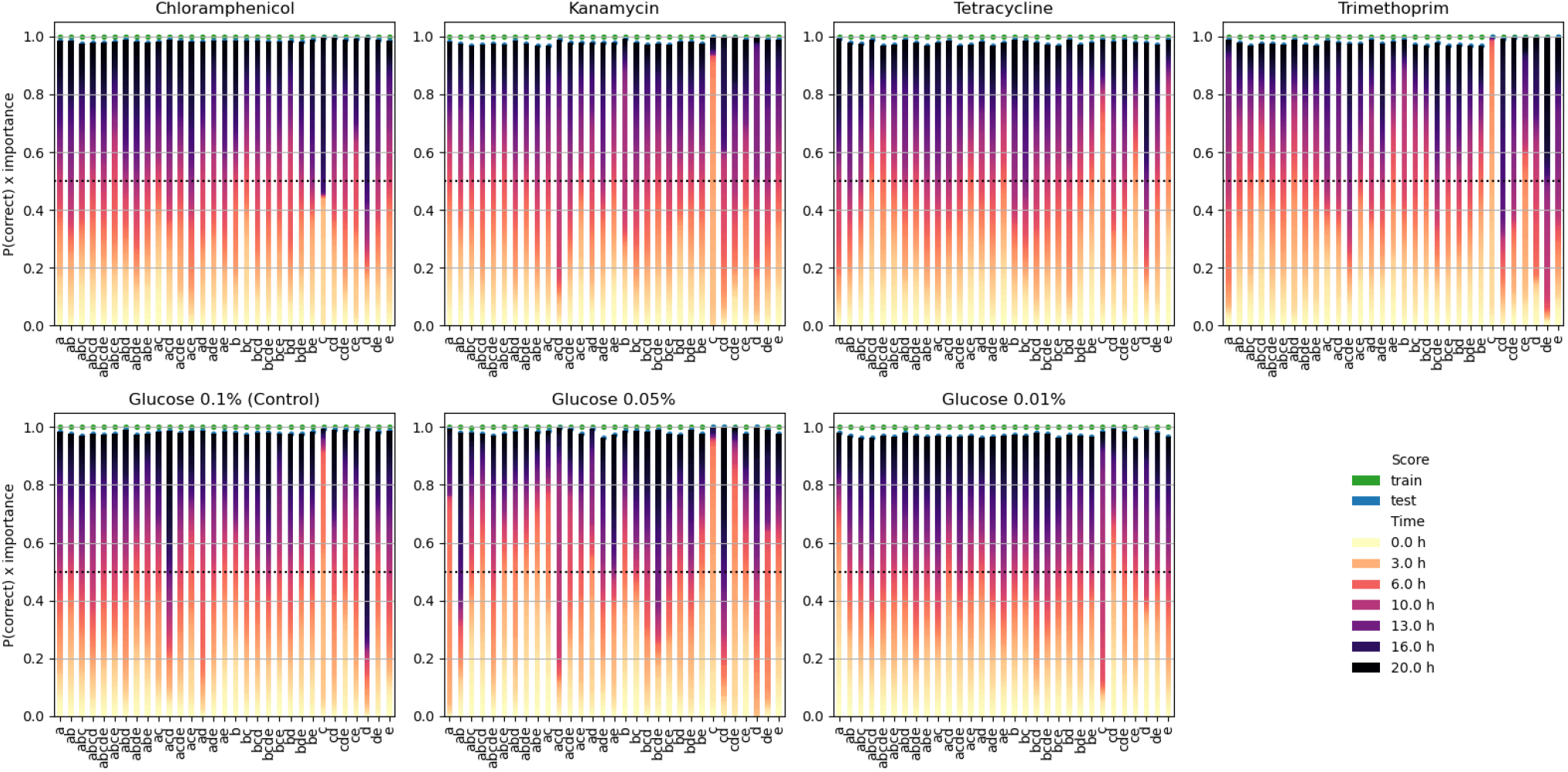
Single-label classification of growth curves. Each population growth curve is to be assessed whether it can be classified for its correspondence to the growth patterns of a given community composition. Each panel represents an environment, for which the classifications take place. The bars are coloured according to the importance of a given time for predicting the correct label, i.e., the shorter the dark part of a bar is, the more growth information is carried by the initial stages of growth.

**Fig S3.**
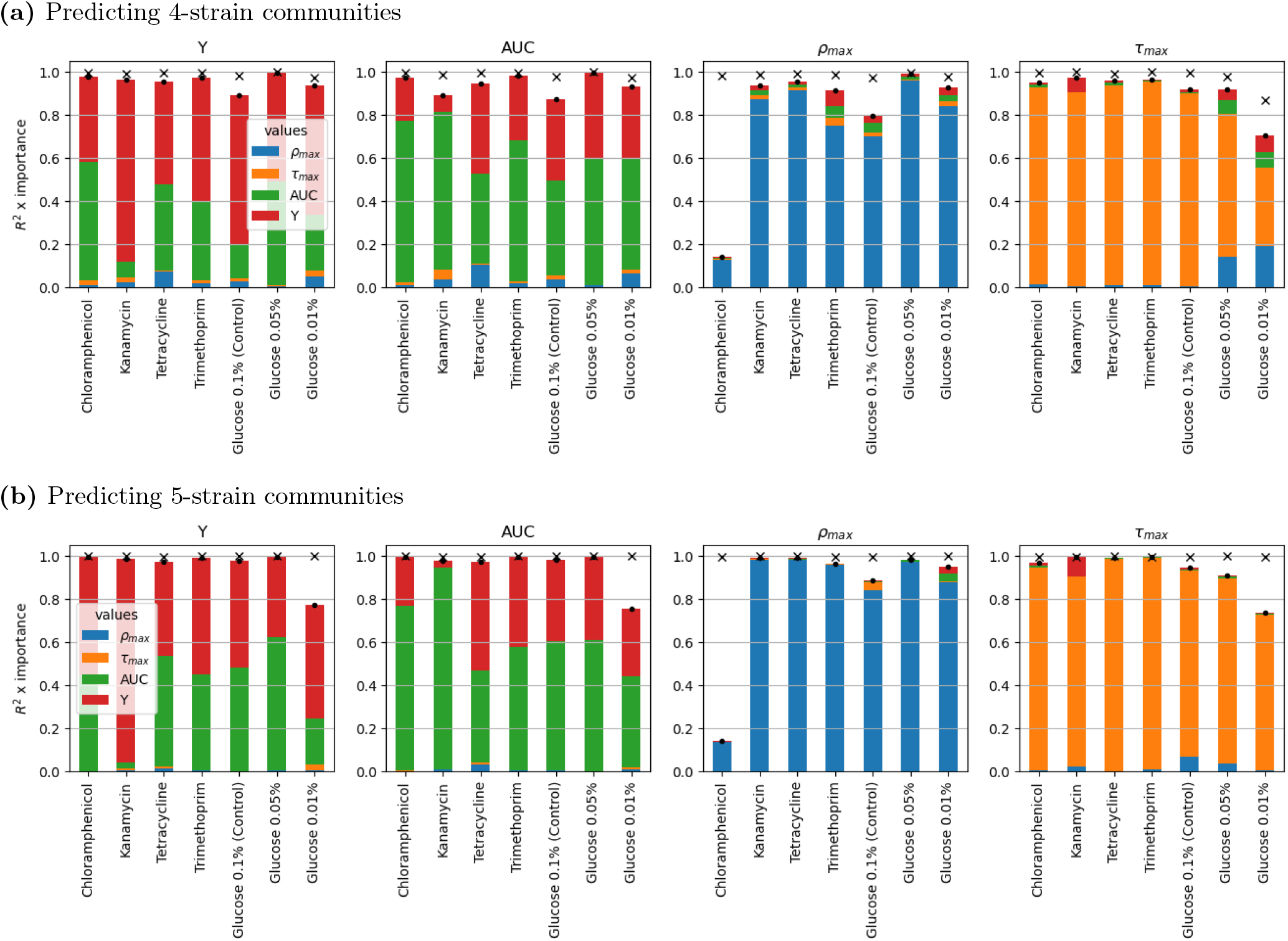
Predicting 4-strain and 5-strain communities from monocultures. A random forest is trained to predict summary values for the shape of the growth curves of 4-strain and 5-strain communities, using the summary values of the involved monocultures as input. Each panel represents the prediction scores and importance of input features for a specific growth summary variable. Each model is trained on train data, which is three quarters of the data of a specific environment, and then scored (*R*^2^) on the train data and test data, which are displayed respectively as a cross and a dot. Each bar then represents the importances of the input features of a random forest, grouped by growth summary variables. These importances are multiplied by the test score.

**Fig S4.**
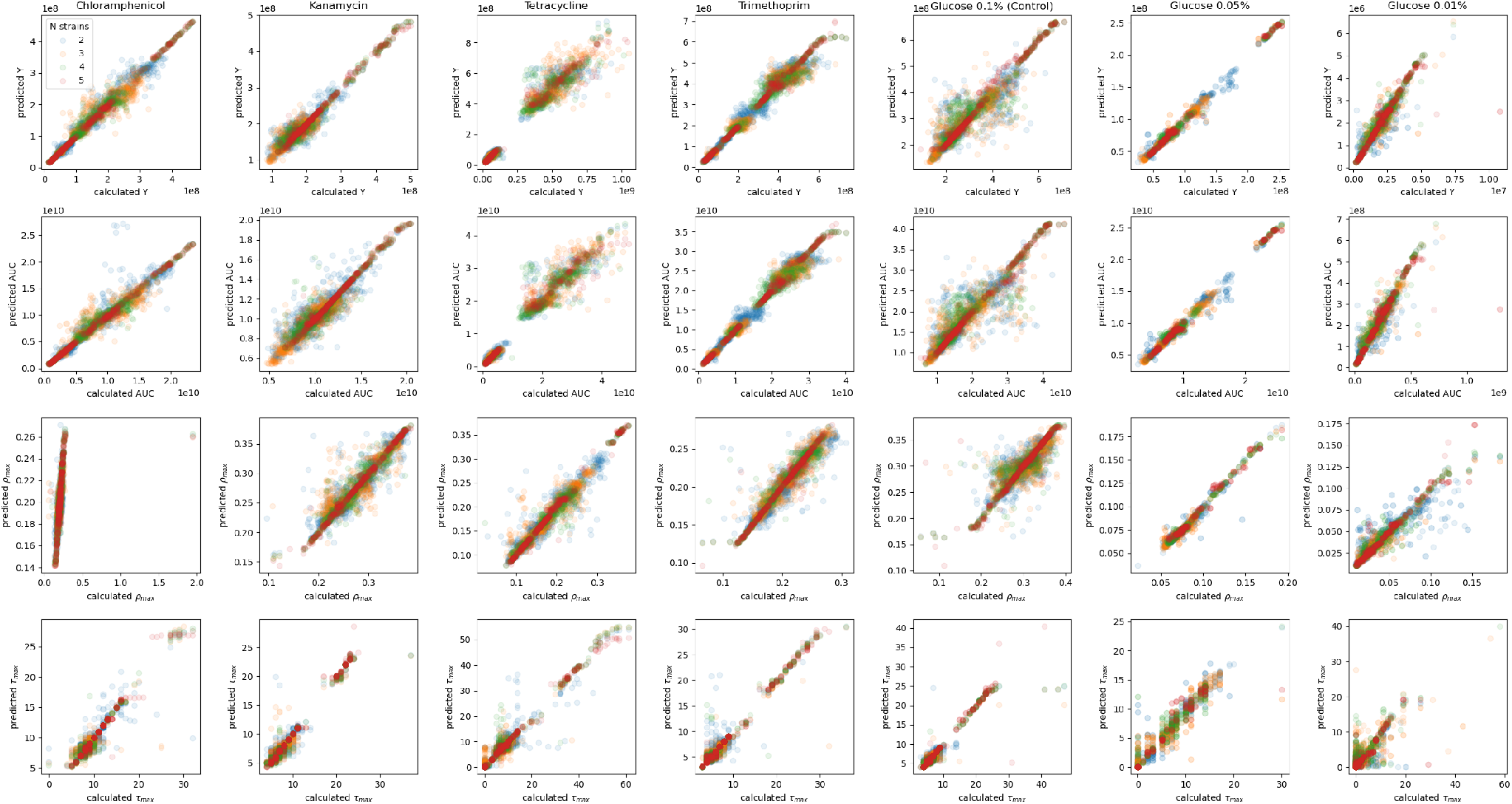
Regressions from 1-strain communities to N-strain communities. Each plot focuses on a given environment and a given curve shape summary value, and compares the values calculated from the experimental data (x axis) to the values predicted by random forests — a good prediction is located at the diagonal of the figure. Each random forest has been trained to predict a given curve shape value for a community combining an arbitrary number N of strains, from all the shape values of the monocultures involved in an N-strain community. The comparisons are coloured according to the number of strains present in the predicted communities. As there is only one way to combine all strains (*N* = 5), the regression task is significantly easier, and the results are good, as expected.

**Fig S5.**
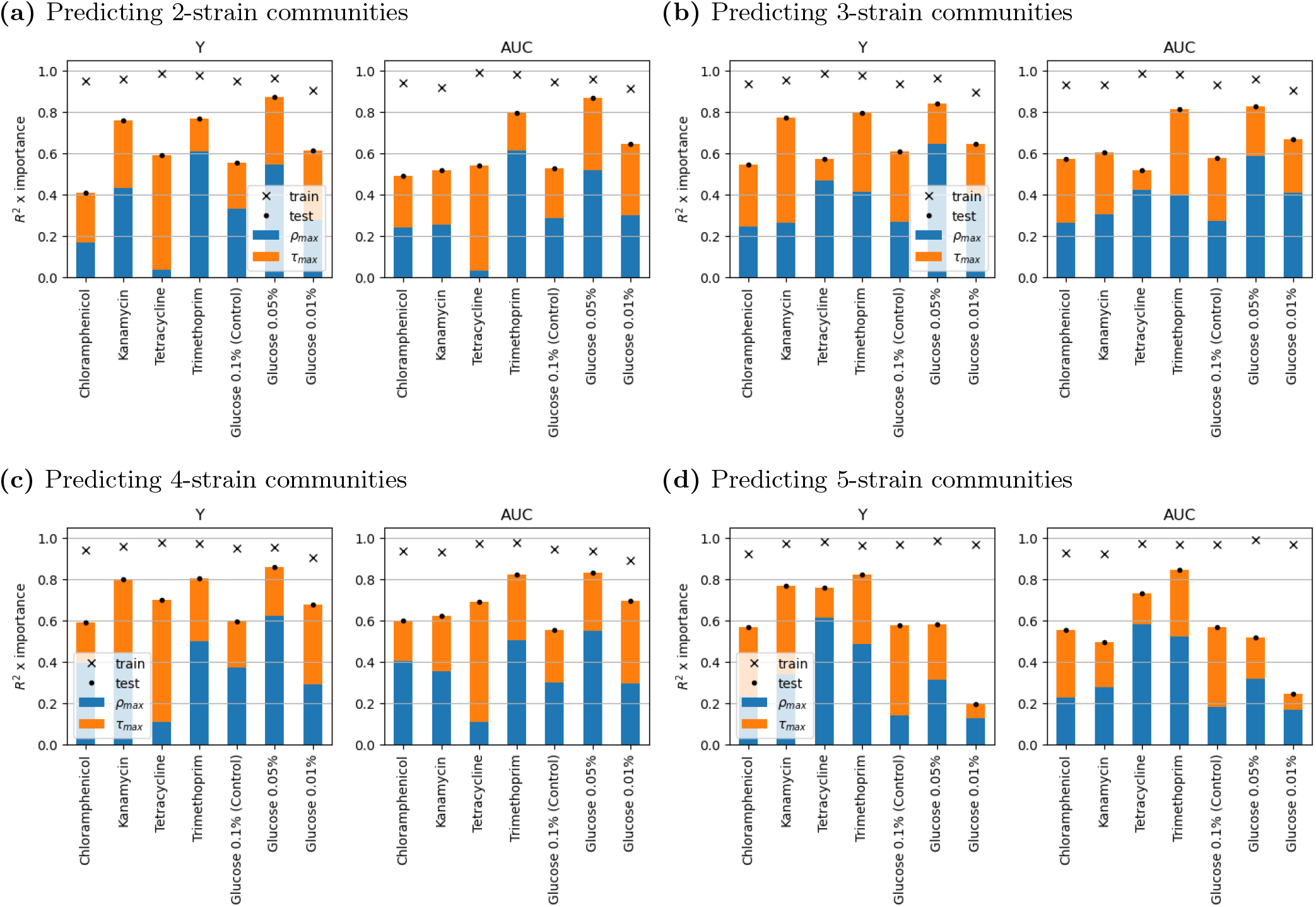
Predicting multi-strain communities from only monoculture *ρ*_max_ and *τ*_max_ values. A random forest is trained to predict summary values for the shape of the growth curves of communities with an arbitrary number of strains, using the maximum growth rate *ρ*_max_ and its timing *τ*_max_ of the involved monocultures as input. Each panel represents the prediction scores and importance of input features for a specific growth summary variable, respectively the final population size *Y* and the shape-independent area under the growth curve (AUC). Each model is trained on train data, which is three quarters of the data of a specific environment, and then scored (*R*^2^) on the train data and test data, which are displayed respectively as a cross and a dot. Each bar then represents the respective importances of *ρ*_max_ and *τ*_max_. These importances are multiplied by the test score.

